# Applying CRISPR-based genetic screens to identify drivers of tumor-cell sensitivity towards NK-cell attack

**DOI:** 10.1101/531962

**Authors:** Klara Klein, Tim Wang, Eric S. Lander, Marcus Altfeld, Wilfredo F. Garcia-Beltran

## Abstract

Natural killer (NK) cells distinguish cancer cells from healthy cells using an array of germline-encoded receptors that interact with ligands expressed on target cells. A balance of inhibitory and activating signals transduced by these receptors regulate NK cell function to provide anti-tumor immunity while maintaining self-tolerance. However, knowledge of the spectrum of factors regulating NK-cell-mediated cytotoxicity, including the contribution of specific ligands and regulatory mechanisms for their expression on tumor cells, remains incomplete. Here, we apply a genome-wide loss-of-function screen in tumor cells using CRISPR/Cas9 technology to identify the factors that promote NK-cell cytotoxicity towards tumor cells. We established the drivers of <u>tu</u>mor-cell <u>se</u>nsitivity towards <u>NK</u>-cell <u>a</u>ttack (TuSeNKA) screening approach using the chronic myeloid leukemia (CML) cell line, K562. Interestingly, we identified B7H6, the ligand for the activating NK cell receptor NKp30, as the single factor whose loss resulted in increased resistance of K562 cells towards NK cells. Our study shows that combination of CRISPR-based genetic screens with NK-cell cytotoxicity assays is a valuable tool for identifying functionally relevant NK cell-tumor cell interactions, paving the way for further investigations that unravel the complexity of signals that promote NK-cell recognition of transformed cells and develop therapies that target these modes of tumor-cell killing.

## INTRODUCTION

NK cells are cytotoxic lymphocytes of the innate immune system and play a critical role in anti-viral and anti-tumor responses. NK cells can engage in direct cytotoxic responses through delivery of granzyme and perforin or via the death receptor pathway, and can also produce large amounts of inflammatory cytokines such as IFN-γ (Smyth *et al.* 2002, Topham and Hewitt 2009, Campbell and Hasegawa 2013). To recognize infected or transformed cells while maintaining self-tolerance to healthy cells, NK cells express a multitude of activating and inhibitory receptors. Inhibitory receptors typically recognize the absence of MHC class I molecules normally expressed on healthy cells (“missing-self” recognition) whereas activating receptors recognize ligands upregulated in cells that have been pathologically altered by stress, infection, or transformation (“induced-self” recognition) (Vivier and Ugolini 2011). NK-cell activity is highly regulated and dependent on integration of signals emanating from both inhibitory and activating receptors, as well as interaction with other immune cells (Diefenbach and Raulet 2001, Raulet and Vance 2006, Lanier 2008, Pegram *et al.* 2011).

Due to their potent activity without the restriction to activation by a specific antigen, NK cells are of growing interest for new immunotherapeutic approaches in cancer (Guillerey *et al.* 2016). For instance, acute myeloid leukemia (AML) patients have shown to benefit from an anti-leukemic effect of NK-cell alloreactivity upon haploidentical hematopoietic stem cell transplantation (Carlsten and Childs 2015, Childs and Carlsten 2015, Guillerey *et al.* 2016, Morvan and Lanier 2016). Furthermore, their cytotoxic potential is currently being exploited in clinical trials that employ inhibitory NK-cell receptor blockade (Benson *et al.* 2012) and *ex-vivo* NK-cell expansion protocols for treatment of cancer patients (Sakamoto *et al.* 2015; Ciurea *et al.* 2017; Ishikawa *et al.* 2018). However, the set of factors regulating NK-cell-mediated cytotoxicity, including the multitude of ligands and regulatory mechanisms for their expression on tumor cells, are far from being completely resolved. Therefore, further research is needed to contribute to a better understanding of factors involved in promoting sensitivity or resistance of tumor cells to NK-cell cytotoxicity.

Genetic screens have elucidated factors involved in interaction of tumor and immune cells (Bellucci *et al.* 2012, Zhou *et al.* 2014, Khandelwal *et al.* 2015, Wucherpfennig and Cartwright 2016, Patel *et al.* 2017). Here, we performed an unbiased CRISPR/Cas9-based loss-of-function screen to investigate functional interactions of primary human NK cells with tumor cells using the well-described NK cell-sensitive chronic myeloid leukemia (CML) cell line K562 (Lozzio and Lozzio 1975, Lozzio and Lozzio 1979, Byrd *et al.* 2007, Brandt *et al.* 2009, Bae *et al.* 2012). We identified B7H6, the ligand for the activating NK-cell receptor NKp30, as the sole factor whose loss resulted in increased resistance of K562 cells towards NK cells, implicating NKp30-mediated killing as the dominating mechanism of NK-cell attack of K562 cells. Altogether, our drivers of <u>tu</u>mor-cell <u>se</u>nsitivity towards <u>NK</u>-cell <u>a</u>ttack (TuSeNKA) screening approach allows for identification of functional interaction of tumor cells with NK cells, which promote sensitivity towards NK-cell-mediated tumor surveillance.

## RESULTS

### The NKp30 ligand B7H6 is a dominant factor in NK-cell-mediated killing of K562 identified in a genome-wide CRISPR/Cas9-based loss-of-function screen

To investigate genes involved in promoting NK-cell activity towards tumor cells (i.e. genes whose loss would increase resistance to NK-cell–mediated killing), we chose K562 as the target cell line. K562 cells lack MHC class I expression and express various activating NK-cell–receptor ligands (including the NKG2D ligands ULBP1,-2 and MIC-A,-B), which in combination engender a high susceptibility to NK-cell–mediated attack (Lozzio and Lozzio 1975, Lozzio and Lozzio 1979, Byrd *et al.* 2007, Brandt *et al.* 2009, Bae *et al.* 2012). We infected Cas9-expressing K562 cells with a genome-wide lentiviral single-guide RNA (sgRNA) library (Wang *et al.* 2015) to generate a pool of K562 cells with individual gene knockouts (**Figure 1A**). Upon selection, 5 × 10^7^ K562 cells were incubated overnight with IL-2-expanded human donor NK cells at a 1:1 effector-to-target (E:T) ratio, which was previously determined to achieve efficient killing of wild-type K562 cells (data not shown). NK cells were then eliminated by IL-2 deprivation and addition of puromycin (to which library-infected K562 cells were resistant) and the surviving K562 cells were expanded under puromycin selection. The surviving K562 population was subjected to a second NK-cell challenge to eliminate cells that might have escaped the initial round of NK-cell killing by chance. sgRNA barcodes from the final, surviving population and an initial cell seeding were quantified by high-throughput sequencing. This allowed the calculation of a “CRISPR gene score”, defined as the average log2 fold-change in the abundance of the five highest scoring sgRNAs, for each gene. Strikingly, our results revealed enrichment of sgRNAs targeting *NCR3LG1* as the single hit from the screen (**Figure 1B**). The surviving K562 population was also sequenced after the first NK-cell challenge before the re-challenge, which similarly revealed loss of *NCR3LG1* as the single prominent hit (data not shown).

**Figure 1.**
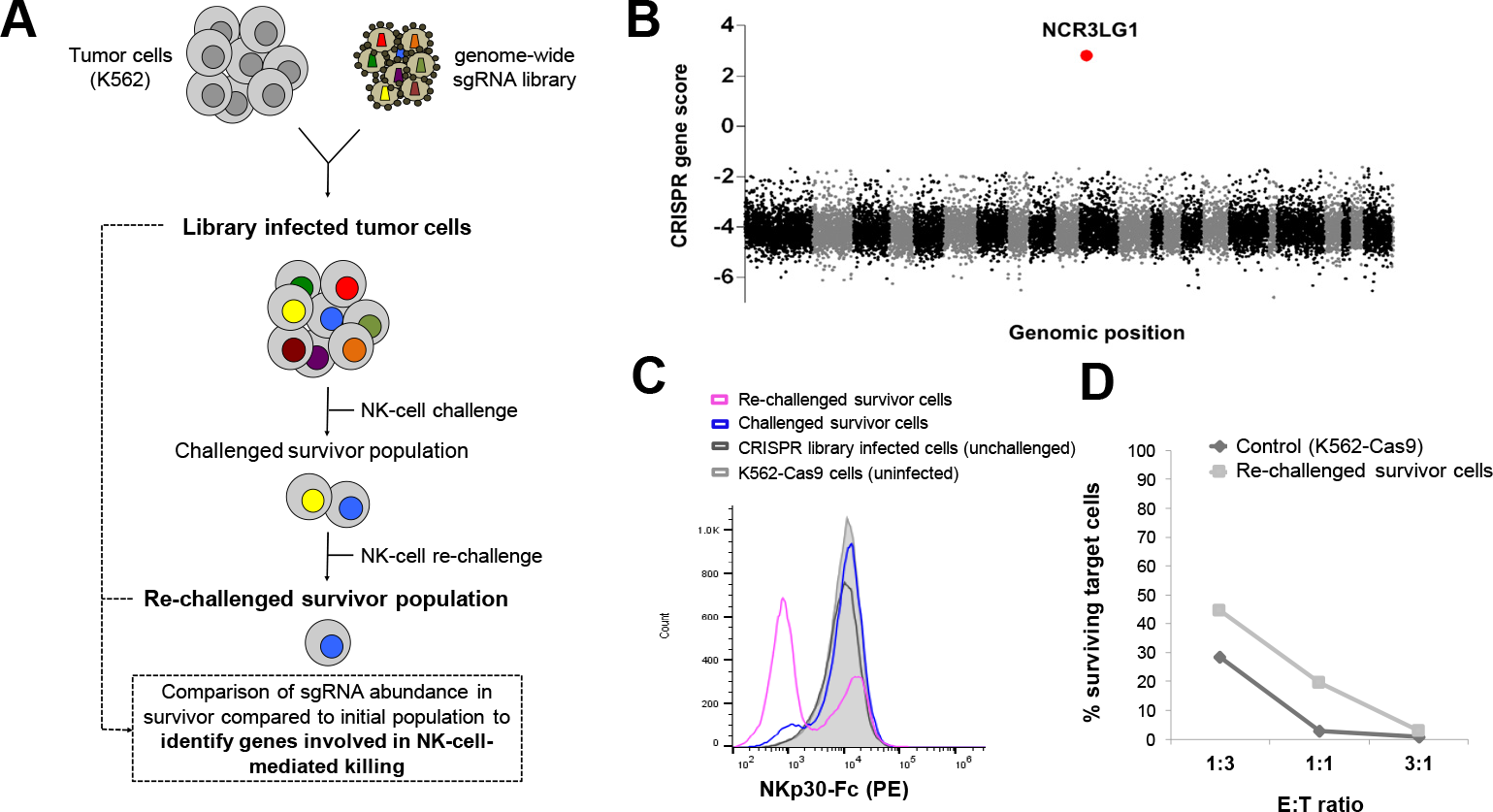
CRISPR/Cas9-based loss-of-function screen identifies *NCR3LG1*, the gene encoding B7H6, as a dominant factor important for NK cell-mediated killing of K562 cells. (A) The screen setup is depicted. 10*10^7^ Cas9-expressing K562 cells (K562-Cas9), co-expressing GFP, were infected with a genome-wide lentiviral sgRNA library to generate a pool of K562 cells with single gene knockouts. The infected cell population was selected with 1μg/mL puromycin for 7 days. 5*10^7^ cells were challenged with IL-2 activated human NK cells at a effector-to-target (E:T) ratio of 1:1 overnight. The challenged survivor population was harvested and expanded in medium containing puromycin to kill off remaining NK cells. A second (re-)challenge of the survivor population was performed under the same challenging conditions. The final survivor population was sequenced to determine sgRNA barcodes and compare abundance in the final compared to the initial unchallenged population. (B) CRISPR gene scores were calculated from sequencing of sgRNA barcodes (as described in Material and Methods) and are depicted for all sgRNA-targeted genes according to their genomic position. Odd (and sex) chromosomes are colored in black and even chromosomes are colored in gray. *NCR3LG1* as the top gene showing sgRNA enrichment in the re-challenged K562 population is indicated in red. (C) The different screen populations were stained with 10 μg/mL of NKp30-Fc contruct (R&D) and PE-coupled secondary antibody and analyzed by flow cytometry. (D) Re-challenged survivor cells and uninfected control K562-Cas9 cells were incubated with IL-2-activated human NK cells at different E:T ratios (1:3, 1:1, 3:1) for 14h. Percentage of surviving target cells, identified as living GFP+ cells, were determined by flow cytometry.

*NCR3LG1* encodes B7H6, a B7 family member and previously discovered ligand for the natural cytotoxicity receptor (NCR) NKp30 (Brandt *et al.* 2009). The dramatic enrichment of cells bearing sgRNAs targeting *NCR3LG1* in the challenged and re-challenged survivors was validated by staining with a soluble NKp30 construct consisting of the extracellular domain of NKp30 fused to Fc region of human IgG1 (NKp30-Fc) (**Figure 1C**). Furthermore, the re-challenged survivor population was killed less efficiently compared to control cells in a dose-dependent manner (**Figure 1D**).

To further validate that loss of B7H6 diminishes the sensitivity of K562 cells to NK-cell killing, we generated *NCR3LG1* knockout cells (B7H6 KO) using individual sgRNAs. Knockout efficiency was validated by flow cytometric analysis with anti-B7H6 antibody staining (**Figure 2A**). Lack of B7H6 expression decreased sensitivity of K562 cells to NK-cell–mediated killing, resulting in an increased survival of B7H6 KO as compared to control cells after 4 and 18 hours of co-incubation with NK cells (**Figure 2B**). This result confirms previous reports showing reduction of NK-cell cytotoxicity towards target cells with decreased B7H6 expression (Cao *et al.* 2015).

**Figure 2.**
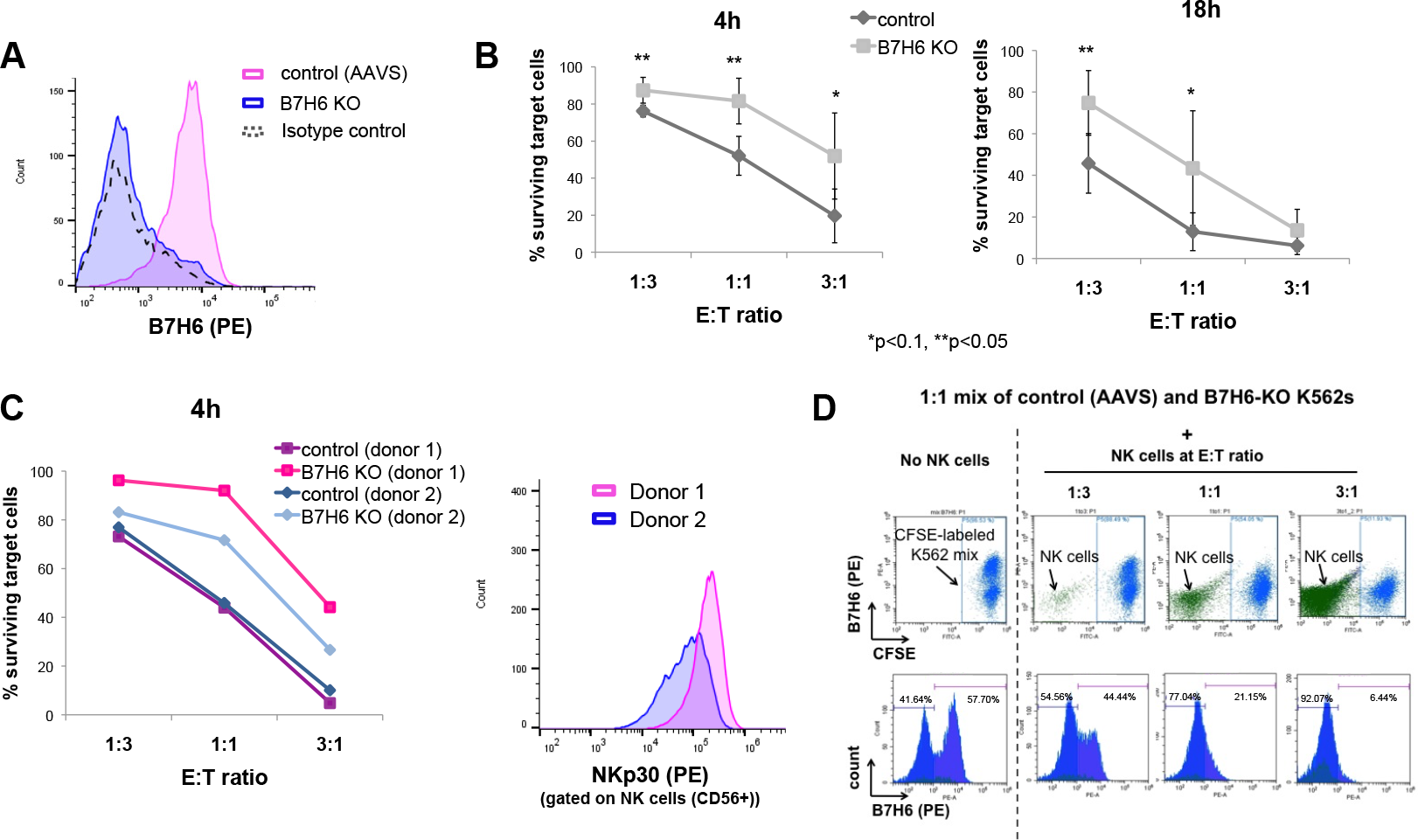
Loss of B7H6 expression decreases sensitivity of K562 cells to NK-cell–mediated cytotoxicity. Individual B7H6 gene knockout (KO) cells were generated in Cas9-expressing K562 cells (K562-Cas9) by infection with lentivirus encoding an sgRNA targeting NCR3LG1/B7H6, while control cells were infected with lentivirus encoding an sgRNA targeting the safe harbour locus AAVS1 as an irrelevant targeting control. (A) Efficiency of sgRNA-mediated B7H6 knockout was confirmed by loss of B7H6 surface expression by flow cytometry compared to control cells. Isotype control for B7H6 staining is shown with a dashed line. (B) Control and B7H6 KO cells were stained with CFSE and incubated at various E:T ratios with IL-2 activated human NK cells for 4h (left panel) or 18h (right panel). Percentage of surviving target cells, determined as living CFSE^+^ cells by flow cytometry, are depicted. Symbols represent the mean with standard deviation of three independent experimental replicates (with different donor NK cells). Significance of differences between survival of control compared to B7H6 KO cells at each E:T ratio were determined by Student’s t-test. *p<0.1, **p<0.05. (C) The graph depicts individual results of cytotoxicity assays with two different donor NK cells (donor 1 and donor 2). Control and B7H6 KO cells were incubated either with donor 1 NK cells (pink) or donor 2 NK cells (blue) at different E:T ratios for 4h. Percentage of surviving target cells were determined by flow cytometric analysis (left panel). NKp30 expression levels on the donor NK cells (gated on CD56^+^ cells) were assessed by flow cytometry (right panel). (D) A 1:1 mix of control and B7H6 KO cells were stained with CFSE and incubated with IL-2 activated human NK cells at different E:T ratios for 4h. After incubation, cells were harvested, washed and stained with PE coupled anti-B7H6 antibody. Dot plots for CFSE fluorescence compared to B7H6-PE fluorescence (upper panel) and histograms for B7H6 fluorescence (lower panel) are shown for the target cell mix only and after incubation with NK cells (CFSE and B7H6 negative cells).

Interestingly, we observed NK cell donor-dependent differences in the efficiency of killing of B7H6 KO compared to control cells (**Figure 2C**). These differences correlated with the level of NKp30 expression on effector NK cells, where NK cells expressing higher NKp30 levels (donor 1) were less efficient in killing of B7H6 KO cells compared to NK cells with lower NKp30 levels (donor 2). This finding might suggest that for NCRs, as previously described for Killer-cell immunoglobulin-like receptors (KIRs), receptor repertoires and expression levels might vary between individuals influenced by genetic and environmental conditions (Pegram *et al.* 2011, Hudspeth *et al.* 2013). Specific NK-cell subsets equipped with specific NK-cell receptor repertoires might therefore be specialized for a specific mode of killing.

Lack of B7H6 surface expression did not confer complete resistance to attack by NK cells, likely due to other activating NK cell ligands and/or lack of MHC class I expression, which can also trigger NK-cell activation. We aimed to confirm that B7H6 KO cells, despite the partially remaining sensitivity to NK-cell cytotoxicity, have a survival advantage over cells that express B7H6 upon NK-cell co-culture, explaining the enrichment of K562s that lost B7H6 expression in the screen. Therefore, we performed a competition cytotoxicity assay, where a 1:1 mix of B7H6 KO and control cells were incubated with NK cells at different E:T ratios over 4 hours (**Figure 2D**). We observed that control target cells (B7H6-PE positive) were preferentially killed compared to B7H6 KO (B7H6-PE negative) cells by NK cells, as indicated by a decrease in the fraction of B7H6-expressing control compared to B7H6 KO cells (**Figure 2D**).

Taken together, performing an unbiased CRISPR-based loss-of-function screen we could identify and further validate B7H6 as an important player in NK-cell–mediated killing of the CML cell line K562. On the basis that its sole loss was sufficient in conferring a partial resistance and survival advantage of K562s in the cytotoxicity screen, B7H6 might be the dominant driver of NK-cell–mediated killing of K562 cells.

## DISCUSSION

Cell-to-cell interactions, many potentially involving unidentified receptor-ligand pairs, play a key role in mediating NK-cell cytotoxicity towards transformed cells. In this study, we describe a drivers of <u>tu</u>mor-cell <u>se</u>nsitivity towards <u>NK</u>-cell <u>a</u>ttack (TuSeNKA) screening approach using CRISPR-based genetic screens to identify key factors promoting NK-cell-mediated tumor-cell killing *in vitro*. Using a genome-wide screening approach, we identified the NKp30 ligand B7H6 as the only gene whose loss resulted in increased survival of cells upon NK-cell challenge. This striking result makes evident that B7H6 is a dominant mediator of NK-cell cytotoxicity towards K562 that was previously underappreciated. Other regulatory factors and NK-cell–receptor ligands are known to contribute to NK-cell killing of K562 cells. However, these genes were not detected in our screen either because of the expression of functionally overlapping ligands (e.g. in case of multiple ligands expressed for the activating NKG2D receptor) or because the stringent conditions employed in our experiment were not sufficiently sensitive to uncover the relatively weaker effects of other potential hits.

There have been several attempt to target NKp30:B7H6 interactions for therapeutic intervention. Bispecific molecules have been described, linking CD20^+^ lymphoma cells to NKp30-expressing NK cells to promote cytotoxic responses. This approach emphasizes that potent NK cell ligands can be exploited to direct specific NK-cell attack towards tumor cells that might not express that ligand themselves (Kellner *et al.* 2012). Furthermore, this potent NK cell–target cell interaction has been adopted for development of B7H6-specific chimeric antigen receptors (CARs) for T-cell based immunotherapy approaches against various tumors known to express B7H6 (Wu *et al.* 2015).

Furthermore, our TuSeNKA screening approach will be a useful tool to determine dominating pathways promoting NK-cell cytotoxicity towards other target cells with less well-defined interaction partners for NK-cell receptors and potentially identify novel factors involved in promoting NK-cell sensitivity. In a first attempt to apply our screening approach to other cell lines, we performed a screen in the AML cell line, SKM1 (Figure S1). However this study did not reveal a clear hit (Figure S1). We posited that, because SKM1 cells display much lower NK-cell sensitivity as compared to K562 cells, many cells carrying irrelevant sgRNAs were able to survive the NK-cell challenge. Interestingly, among the top-scoring genes were several pro-apoptotic molecules (e.g. *PMAIP1, BID*) and the TRAIL receptor (*TNFRSF10A*) (Figure S1), implying that death-receptor signaling might play a major role in killing of the AML cell line SKM1.

CRISPR-based screens studying tumor progression and metastasis have been previously performed in an *in-vivo* setting (Chen *et al.* 2015). It might be interesting to perform screens to study the interaction between CRISPR library-infected tumor cells and NK cells upon transplantation into mice, lacking the adaptive immune system, to evaluate dominant factors promoting NK-cell cytotoxicity towards specific tumor cells *in vivo*. Also, similar to previous *in vivo* shRNA screens in T cells (Zhou *et al.* 2014), factors involved in regulating NK-cell function *in vivo* and factors promoting NK-cell tumor infiltration might be studied utilizing the CRISPR/Cas9 system. However, these investigations are complicated as primary NK cell are difficult to transduce. Therefore, optimization of protocols for genetic manipulation of primary NK cells are crucial. Alternatively, these complications might be bypassed by lentiviral infection of hematopoietic stem cells in combination with *in vitro* differentiation protocols or generation of NK cells upon transplantation into mice (Carlsten and Childs 2015, Childs and Carlsten 2015, Wucherpfennig and Cartwright 2016).

Overall, we demonstrate that the CRISPR/Cas9 system and CRISPR-based genetic screens can serve as powerful tools to study questions related to tumor immunology. However, there are still some limitations to CRISPR-based genetic screening approaches (Morgens *et al.* 2016). First, functionally overlapping genes cannot be studied using single sgRNA libraries. Second, the effects of essential genes cannot be investigated in a positive selection screen, as loss of these genes will cause cells to be eliminated from the population. Additionally, pooled genetic screening approaches are not amenable for probing cell-non-autonomous processes such as the actions of secreted factors. These current limitations of CRISPR-based screens can be overcome by further optimization and supplementation with other technical approaches. Altogether, these combined CRISPR-based functional screening approaches will help to identify dominant and functionally relevant receptor-ligand interactions among multiple interactions, uncovering key therapeutic targets and allows us to address unanswered questions in tumor biology and immunology (Sanchez-Rivera and Jacks 2015, Wucherpfennig and Cartwright 2016).

## MATERIAL AND METHODS

### Cell culture and human samples

All tumor cell lines, including transductants, were cultured in IMDM (Gibco) supplemented with 20% of heat-inactivated FBS, 100 U/mL penicillin, 100 μg/mL streptomycin and 2 mM L-glutamine (all reagents from Gibco) at 37°C/5% CO_2_. Primary human NK cells were isolated from leukocyte-enriched peripheral blood of healthy human donors using the RosetteSep™ Human NK Cell Enrichment Cocktail (StemCell Technologies) according to manufacturer’s instructions. Primary human NK cells were cultured in RPMI supplemented with 10% heat-inactivated FBS, 100 U/mL penicillin, 100 μg/mL streptomycin and 2 mM L-glutamine (all reagents from Gibco) in the presence of 100 U/mL IL-2 (Peprotech) at 37°C/5% CO_2_. Leukocyte-enriched peripheral blood was obtained from healthy human donors who had provided informed consent and was used in accordance to protocols approved by Partners Human Research Committee and Institutional Review Board of Massachusetts General Hospital.

### Cloning and lentiviral transduction of individual sgRNAs and Cas9 constructs

Individual sgRNA constructs targeting NCR3LG1/B7H6 (sgNCR3LG1 sequences: GGGTGACCACCACCTCACAT and AACTCCTCTCAGGAAGACCC) and AAVS1 (sgAAVS1: GGGGCCACTAGGGACAGGAT) were cloned into lentiCRISPR-v1 (Addgene) or sgOpti (Fulco *et al.* 2016), a modification of pLenti-sgRNA vector (Addgene #71409) with an optimized sgRNA scaffold (Chen *et al.* 2013), as described previously (Cong *et al.*, 2013; Wang *et al.* 2016). For lentivirus production, HEK293T cells were transfected with the respective sgRNA-containing plasmids together with the VSV-G (pCMV-VSV-G) envelope plasmid and dVPR (pCMV-dR8.2) packaging plasmids (from Addgene) using the XtremeGene9 transfection reagent (Roche). Media change was performed after overnight incubation and viral supernatant was harvested after additional 36h. Target cells were infected in media containing 8 μg/mL of polybrene (EMD Millipore) by centrifugation at 2220 RPM for 45 min. After overnight incubation, remaining virus was removed by centrifugation. Selection of infected cells was performed with puromycin (Gibco) starting 24-48 h after media change. For generation of stable Cas9-GFP expressing cell lines, cells were transduced with a lentiviral construct expressing Cas9-2A-GFP, a version of lentiCRISPR-v1 in which the puromycin N-acetyltransferase ORF was replaced with eGFP (Wang *et al.* 2017). Cas9-GFP positive cells were sorted out of the transduced bulk cell population by FACS and expanded for subsequent transduction of individual sgRNAs.

### Lentiviral sgRNA library construction

The genome-scale lentiviral sgRNA library was designed and generated by Tim Wang as previously described (Wang *et al.* 2015), including 178,896 sgRNAs targeting 18,166 protein-coding genes in the human consensus CDS (CCDS) and 1,004 non-targeting control sgRNAs. Briefly, oligonucleotide pools were synthesized on CustomArray 90K arrays for generation of sgRNA library with PCR tags. The library was amplified by PCR with primers adding homology arms for Gibson assembly. The PCR product was purified and assembled into BsmBI (NEB) digested vector backbones in a Gibson reaction (NEB). The product was cleaned up with AMPure XP SPRI beads (Beckman Coulter) and electroporated into Endura competent cells (Lucigen). Cells were expanded in an overnight liquid culture and the pooled plasmid library was extracted using QIAfilter Plasmid Maxi Kit (Qiagen). If necessary, an additional purification step via ethanol precipitation was performed. The obtained pooled plasmid library was used to produce lentiviral sgRNA library as described.

For screens in K562 cells, 1 × 10^8^ Cas9-GFP expressing derivatives were infected with the genome-scale sgRNA library lentivirus at a low MOI.

### NK-cell cytotoxicity assay

Target cells either expressed GFP (K562-Cas9) or were stained with CFSE (Life Technologies). For CFSE staining, target cells were washed with PBS and resuspended at a concentration of 1 × 10^6^ cells/mL in 0.5 μM CFSE in PBS. Cells were stained for 5 min at 37°C and the reaction was stopped by adding RPMI containing 10% FBS. Primary human NK cells were used as effector cells after culture in 100 U/mL IL-2 for at least 3 days. 50.000 target cells were mixed with effector cells at different effector-to-target (E:T) cell ratios (3:1, 1:1, 1:3) and co-incubated in 96-well plates for 4-24 h at 37°C/5% CO_2_. NK cell-mediated killing of target cells was analyzed on a BD Accuri™ C6 Cytometer and depicted as percentage of remaining surviving green-fluorescent cells upon incubation with NK cells from the initial count of living green-fluorescent target cells (cells cultured without effector cells).

### <u>Tu</u>mor-cell <u>se</u>nsitivity towards <u>NK</u>-cell attack (TuSeNKA) screening procedure

After 7 days of selection with 1μg/mL puromycin, 5 × 10^7^ genome-scale lentiviral sgRNA library infected K562-Cas9 cells were harvested and washed with PBS. Cells were mixed with primary human NK cells at E:T ratio of 1:1 and seeded into 96-well plates with 200,000 target cells per well and incubated overnight (16 h) in IL-2 containing RPMI medium. Cells were harvested, washed in PBS to remove IL-2 and surviving cells were cultured in 20% FBS IMDM containing 1 μg/mL puromycin to kill off remaining effector cells. The surviving target cell population was expanded to a sufficient amount of cells for about 1-2 weeks and subjected to a re-challenge with primary NK cells, following the same protocol as described in a smaller scale. The resulting re-challenged survivor cell population was harvested for genomic DNA extraction and subsequent high-throughput sequencing as described below to determine sgRNAs targeting genes involved in NK-cell-mediated killing of K562 target cells.

### Flow cytometric analysis

For flow cytometric analysis, cells were washed twice with PBS and incubated with fluorescently labeled antibodies or isotype controls (Table 1) (at 1:20-1:100 dilutions depending on the antibody) in the presence of human Fc receptor blocking reagent (1:10) (Miltenyi Biotec) for 15 min at 4°C. Unbound antibodies were washed away with PBS, cells were resuspended in PBS and subsequently analyzed on a CytoFLEX Flow Cytometer (Beckman Coulter). For staining with soluble receptor constructs (R&D), cells were first stained with 20 μg/mL of soluble receptor-Fc constructs (Table 1) for 15min at 4°C. After a PBS wash, cells were stained with a PE-conjugated anti-human IgG1(Fc) secondary antibody (1:50) (ThermoFisher) for additional 15min, followed by a PBS wash prior to acquisition. As a negative/background control for soluble receptor construct stainings, cells were stained with the secondary antibody alone (2^nd^ only). Data were analyzed using FlowJo software.

**Table 1:** Antibodies and constructs used

| **Table 1: Antibodies and constructs used** |  |  |  |  |
| --- | --- | --- | --- | --- |
| **Specificity (clone)** | **Fluorophore** | **Isotype** | **Company** | **Catalog No.** |
| anti-human CD56 (HCD56) | Brilliant Violet 421™ | mouse IgG1, κ | BioLegend | 318327 |
| anti-human NKp30 (P30-15) | PE | mouse IgG1, κ | BioLegend | 325207 |
| anti-human B7H6 (#875001) | PE | mouse IgG1 | R&D Systems | AB7144P-025 |
| isotype control for B7H6 (#11711) | PE | mouse IgG1 | R&D Systems | IC002P |
| F(ab')<sub>2</sub> goat anti-human IgG(Fc) (polyclonal) | PE | goat IgG (F(ab)<sub>2</sub>) | ThermoFisher | H10104 |
| recombinant human NKp30-Fc chimera | none (purified) | human IgG1 (Pro100 – Lys330) | R&D Systems | 1849-NK-025 |

### High-throughput sequencing and CRISPR score calculation

Sequencing of sgRNA barcodes and analysis of sgRNA enrichment (CRISRP score) were performed similar as previously described (Wang *et al.* 2014, Wang *et al.* 2015). Briefly, genomic DNA was isolated from sgRNA library infected cell populations using the QIAamp DNA Blood Midi Kit (Qiagen) according to manufacturer’s instructions.

The sgRNA inserts were amplified from the isolated genomic DNA in a PCR reaction using a universal forward primer (5’→3’): ATGATACGGCGACCACCGAGATCTAGAATACTGCCATTTGTCTCAAG and barcoded sample-specific reverse primers (5’→3’): CAAGCAGAAGACGGCATACGAGATC(N)_6_TTTCTTGGGTAGTTTGCAGTTTT where (N)_6_ denotes the sample barcode.

The resultant PCR products were purified, quantified, and sequenced on a MiSeq (Illumina) using the following primers (5’→3’):

Illumina sequencing primer: CGGTGCCACTTTTTCAAGTTGATAACGGACTAGCCTTAT TTTAACTTGCTATTTCTAGCTCTAAAAC
Illumina indexing primer: TTTCAAGTTACGGTAAGCATATGATAGTCCATTTTAAAACATAATTTTAAAACTGCAAACTACCCAAGAAA

Sequencing reads were aligned to the sgRNA library and the abundance of each sgRNA was calculated. Log2-fold changes for each sgRNA at the endpoint (after NK cell (re)-challenge) compared to the initial timepoint (sgRNA abundance at timepoint of initial library infection) were calculated. Gene-based CRISPR scores for the genome-scale screens were defined as the average of these log2-fold changes of the 5 most highly scoring sgRNAs (from the total of 10 sgRNAs per gene).

## ACKNOWLEDGEMENTS

This work was supported by the US National Institute of Health (F31 CA189437 to T.W.; R01-AI067031-08 to M.A.; F31AI116366 to W.F.G.-B.), the National Institute of General Medical Sciences (T32GM007753 to W.F.G.-B.), the National Human Genome Research Institute (2U54HG003067-10 to E.S.L.), and the MIT Whitaker Health Sciences Fund (to T.W.), the Broad Institute of MIT and Harvard, and the Ragon Institute of MGH, MIT and Harvard. E.S.L. directs the Broad Institute, which holds patents and has filed patent applications on technologies related to CRISPR-Cas9. E.S.L., M.A., W.F.G.-B. and K.K. have no personal financial interest in the work in the paper. T.W. is a co-founder of KSQ Therapeutics, which is using CRISPR-based screens to identify drug targets. T.W. and E.S.L. are inventors on a patent for functional genomics using CRISPR-Cas9 (US 15/141,348). The content is solely the responsibility of the authors and does not necessarily represent the official views or policies of the National Institute of General Medical Sciences, the National Institutes of Health, or the Department of Health and Human Services, nor does mention of trade names, commercial products, or organizations imply endorsement by the US Government.

## SUPPLEMENTARY FIGURE

**Figure S1.**
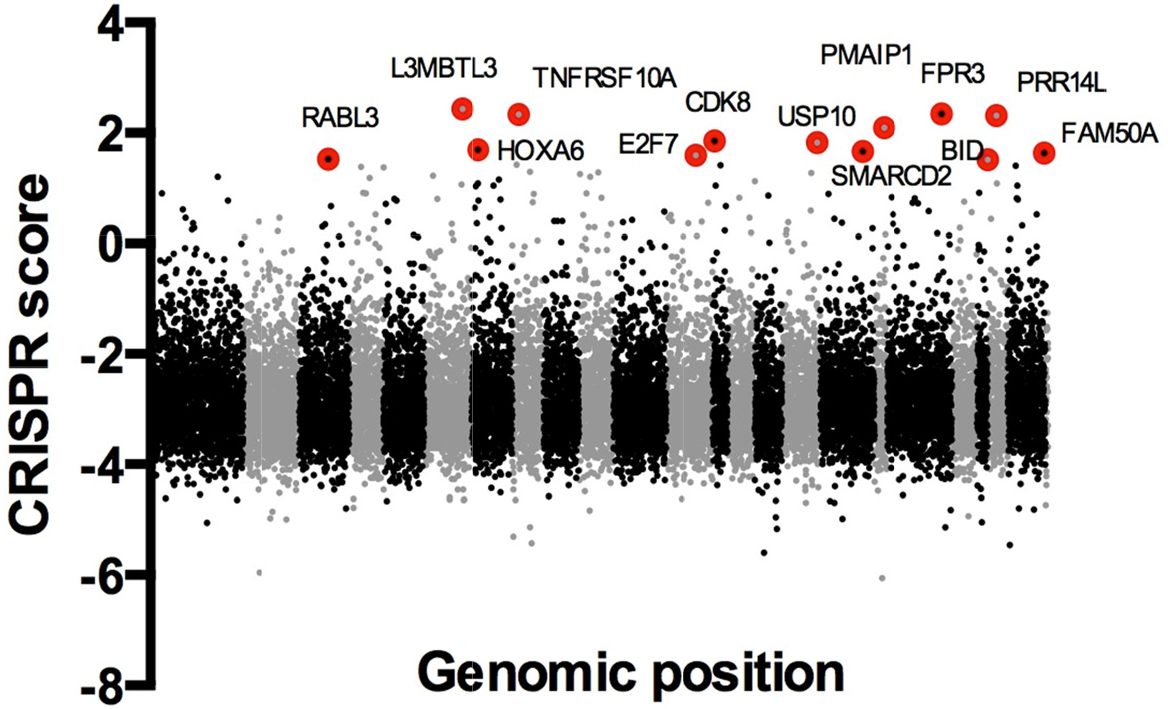
CRISPR/Cas9 screen indicates pathways involving death receptor signaling to play a role in NK cell-mediated killing of SKM1 cells. The screen setup was performed as described for K562 cells (Fig. 1), however 5 × 10^7^ sgRNA library infected Cas9-GFP expressing SKM1 cells were incubated with IL-2 activated human NK cells at a effector-to-target (E:T) ratio of 2:1 overnight. The challenged survivor population was harvested and expanded in medium containing puromycin to kill off remaining NK cells. A second (re-)challenge of the survivor population was performed under the same challenging conditions. The final survivor population was sequenced to determine sgRNA barcodes and compare abundance in the final compared to the initial population. CRISPR gene scores were calculated from sequencing of sgRNA barcodes (as described in Material and Methods) and are depicted for all sgRNA-targeted genes according to their genomic position. Odd (and sex) chromosomes are colored in black and even chromosomes are colored in gray. Genes showing the highest sgRNA enrichment are indicated in red.

